# Specialised functions of two common plasmid mediated toxin-antitoxin systems, *ccdAB* and *pemIK*, in *Enterobacteriaceae*

**DOI:** 10.1101/2020.03.06.980490

**Authors:** Alma Y. Wu, Muhammad Kamruzzaman, Jonathan R. Iredell

## Abstract

Toxin-antitoxin systems (TAS) are commonly found on bacterial plasmids and are involved in plasmid maintenance. Even though the same TAS are present in a variety of plasmid types and bacterial species, differences in their sequences, expression and functions are not well defined. Here, we aimed to identify commonly occurring plasmid TAS in *Escherichia coli* and *Klebsiella pneumoniae* and compare the sequence, expression and plasmid stability function of their variants. 27 putative type II TAS were identified from 1063 plasmids of *Klebsiella pneumoniae* in GenBank. Among these, *ccdAB* and *pemIK* were found to be most common, also occurring in plasmids of *E. coli*. Comparisons of *ccdAB* variants, taken from *E. coli* and *K. pneumoniae*, revealed sequence differences, while *pemIK* variants from IncF and IncL/M plasmids were almost identical. Similarly, the expression and plasmid stability functions of *ccdAB* variants varied according to the host strain and species, whereas the expression and functions of *pemIK* variants were consistent among host strains. The specialised functions of some TAS may determine the host specificity and epidemiology of major antibiotic resistance plasmids.

## Introduction

Toxin-antitoxin systems (TAS) were originally discovered on bacterial plasmids in the 1980s, [1, 2] but have since also been recognised on bacterial chromosomes [3]. TAS cassettes typically consist of two gene loci, governed by a common regulation mechanism [4], encoding a stable toxin that induces cell death or arrests growth, and a labile antitoxin that neutralises the toxin through binding to the toxin or other means. While the toxin is always a protein, the antitoxin can be protein or RNA based, and thus TAS can be categorised into six different types (Types I-VI), based on the nature and mechanism of action of the antitoxin [5, 6]. The Type II system, in which both toxin and antitoxin are proteins, is the typical model of a TAS and the best studied, and is probably the most common in bacteria [7].

It has long been known that TAS play a role in plasmid maintenance through postsegregational killing of plasmid free cells [2, 8]. Within *Enterobacteriaceae*, TAS are common among conjugative plasmids including antibiotic resistance (AbR) plasmids, and often associated with certain plasmid incompatibility (Inc) types. For example, the type II TAS *pemIK* and *vagCD* are usually found on IncF and IncL/M plasmids, and IncF and IncHI2 plasmids respectively [9, 10].

The distribution of TAS in the plasmids found in *Escherichia coli* is well described [11-13], but very little is known about the distribution of TAS in the plasmids residing in *Klebsiella pneumoniae* species. There is also a general assumption that common TAS (such as *ccdAB*) found in different plasmid types or in different bacterial species have similar functions, but functional differences in TAS have previously been noted [14]. For example, the *hok-sok* TAS from plasmid R1 and *E. coli* K-12 chromosome are involved in postsegregational killing and persister cell formation respectively [15, 16].

In this study, we analysed the distribution of type II TAS in 1063 fully sequenced plasmids found in *K. pneumoniae* retrieved from GenBank. Variation in DNA sequence, expression and function of two common TAS found in both *E. coli* and *K. pneumoniae* plasmids, *ccdAB* and *pemIK*, were examined.

*ccdAB* is a well-studied, primarily plasmid-associated type II TAS, although copies have also been found on bacterial chromosomes where it appears to be involved in the bacterial stress response [17, 18]. The toxin, ccdB, binds to and inhibits the action of DNA gyrase, thus inhibiting DNA replication [19]. *pemIK*, another well studied type II TAS that is related to the *mazEF* system, is an mRNA endoribonuclease that inhibits protein synthesis [20]. Here, we provide vital information about the similarity, specificity and functions of these two TAS, with broad implications for their role in the spread of antibiotic resistance.

## Materials and Methods

### Identification of TAS on *K. pneumoniae* plasmids

The names and complete sequences of all plasmids from *K. pneumoniae* species available in GenBank at the time of search (August 2019) were retrieved (https://www.ncbi.nlm.nih.gov/genome/?term=klebsiella+pneumoniae), and the sequences of all plasmids >30 kb examined with TA Finder (http://202.120.12.133/TAfinder/TAfinder.php) [21] using the default parameters to identify potential type II TAS. The plasmid incompatibility (Inc) type of each plasmid was defined using PlasmidFinder (https://cge.cbs.dtu.dk/services/PlasmidFinder/) [22].

### Alignment of TAS sequences

Representative examples of *ccdAB* and *pemIK* TAS were chosen, and the nucleotide and amino acid sequences of the toxin and antitoxin coding regions retrieved from GenBank (https://www.ncbi.nlm.nih.gov/genbank/) using coordinates obtained from TA Finder. These sequences were aligned in MEGA7 (http://www.megasoftware.net/mega7/) [23] using the ClustalW algorithm. The amino acid sequences were then used to predict the secondary structures of the proteins using PsiPred (http://bioinf.cs.ucl.ac.uk/psipred/) [24].

TAS promoters were predicted using BPROM (http://www.softberry.com/berry.phtml?topic=bprom&group=programs&subgroup=gfindb) (Softberry) with default parameters, with the input being the 500 bp region upstream of the ATG start codon of the antitoxin gene. The putative promoter sequences were then aligned as described above.

### Plasmids, bacteria, primers and culture conditions

Tables 1, 2 and 3 list plasmids, bacterial strains and primers respectively. Bacteria were grown in Luria-Bertani (LB) broth (BD Biosciences, NJ, USA), with kanamycin (50 µg/mL) or chloramphenicol (20 µg/mL) (Sigma-Aldrich, MO, USA) added as indicated. Insertion of TAS into vectors was carried out using standard restriction digestion and ligation cloning protocols, and chemical transformation and electroporation into host strains was also performed using standard protocols. The plasmid construct details can be found in Table 1. Each solution used was rendered sterile either through autoclaving or filter sterilising at 0.22 µm.

**Table 1.**
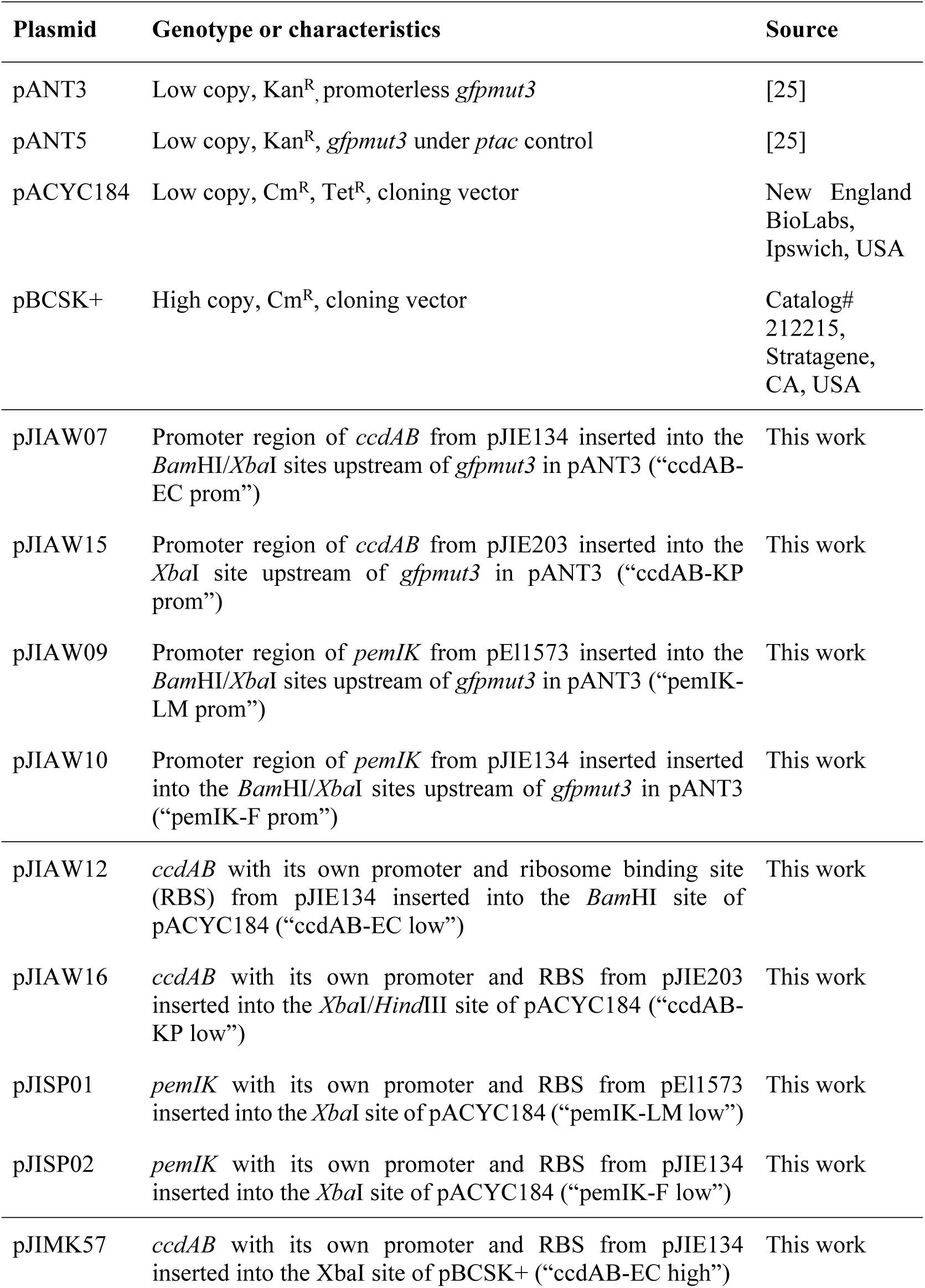
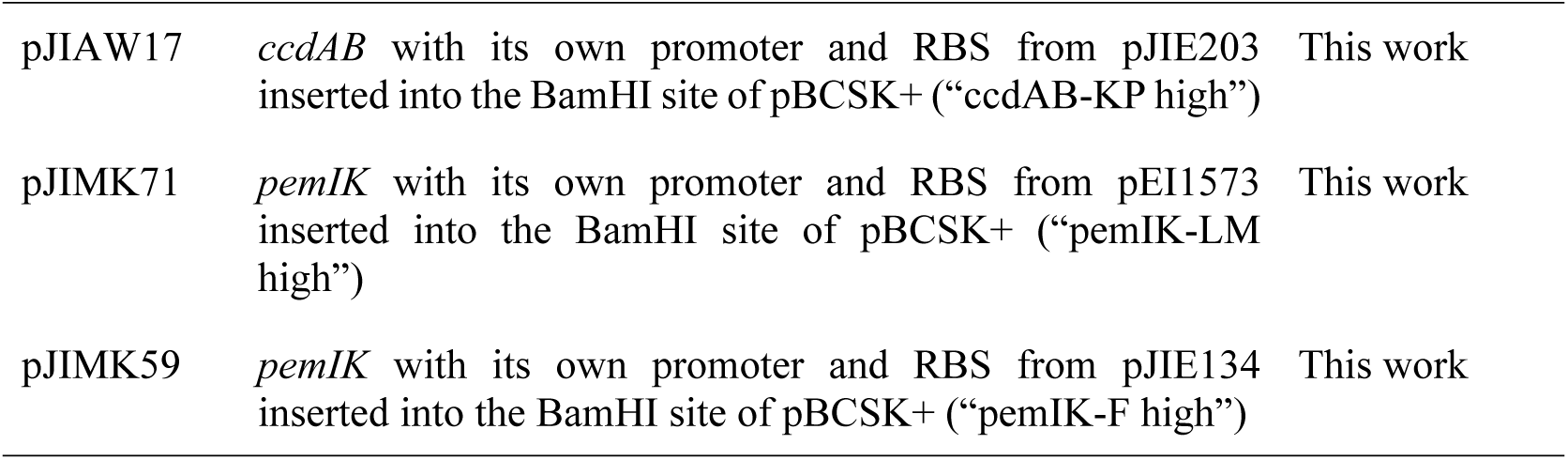
Plasmids used in this study.

| Plasmid | Genotype or characteristics | Source |
| --- | --- | --- |
| pANT3 | Low copy, Kan<sup>R</sup>, promoterless *gfpmut3* | [25] |
| pANT5 | Low copy, Kan<sup>R</sup>, *gfpmut3* under *ptac* control | [25] |
| pACYC184 | Low copy, Cm<sup>R</sup>, Tet<sup>R</sup>, cloning vector | New England BioLabs, Ipswich, USA |
| pBCSK+ | High copy, Cm<sup>R</sup>, cloning vector | Catalog# 212215, Stratagene, CA, USA |
| pJIAW07 | Promoter region of *ccdAB* from pJIE134 inserted into the *Bam*HI/*Xba*I sites upstream of *gfpmut3* in pANT3 (“*ccdAB*-EC prom”) | This work |
| pJIAW15 | Promoter region of *ccdAB* from pJIE203 inserted into the *Xba*I site upstream of *gfpmut3* in pANT3 (“*ccdAB*-KP prom”) | This work |
| pJIAW09 | Promoter region of *pemIK* from pEI1573 inserted into the *Bam*HI/*Xba*I sites upstream of *gfpmut3* in pANT3 (“*pemIK*-LM prom”) | This work |
| pJIAW10 | Promoter region of *pemIK* from pJIE134 inserted inserted into the *Bam*HI/*Xba*I sites upstream of *gfpmut3* in pANT3 (“*pemIK*-F prom”) | This work |
| pJIAW12 | *ccdAB* with its own promoter and ribosome binding site (RBS) from pJIE134 inserted into the *Bam*HI site of pACYC184 (“*ccdAB*-EC low”) | This work |
| pJIAW16 | *ccdAB* with its own promoter and RBS from pJIE203 inserted into the *Xba*I/*Hind*III site of pACYC184 (“*ccdAB*-KP low”) | This work |
| pJISP01 | *pemIK* with its own promoter and RBS from pEI1573 inserted into the *Xba*I site of pACYC184 (“*pemIK*-LM low”) | This work |
| pJISP02 | *pemIK* with its own promoter and RBS from pJIE134 inserted into the *Xba*I site of pACYC184 (“*pemIK*-F low”) | This work |
| pJIMK57 | *ccdAB* with its own promoter and RBS from pJIE134 inserted into the *Xba*I site of pBCSK+ (“*ccdAB*-EC high”) | This work |
| pJIAW17 | *ccdAB* with its own promoter and RBS from pJIE203 inserted into the BamHI site of pBCSK+ (“*ccdAB*-KP high”) | This work |
| pJIMK71 | *pemIK* with its own promoter and RBS from pEI1573 inserted into the BamHI site of pBCSK+ (“*pemIK*-LM high”) | This work |
| pJIMK59 | *pemIK* with its own promoter and RBS from pJIE134 inserted into the BamHI site of pBCSK+ (“*pemIK*-F high”) | This work |

**Table 2.** Bacterial strains used in this study.

| Strain | Genotype or characteristics | Source |
| --- | --- | --- |
| ***E. coli* strains** |  |  |
| DH5α | Host strain used for cloning.<br>F- ϕ80*lacZ*ΔM15<br>$\Delta$(*lacZYA-argF*)U169<br>*recA1 endA1 hsdR17*(r<sub>k</sub><sup>-</sup><br>mk<sup>+</sup>) *phoA supE44 thi-1*<br>*gyrA96 relA1 deoR* | Invitrogen (Carlsbad, USA) |
| Ec WH62 | Antibiotic sensitive clinical<br>*E. coli* isolate | Westmead Hospital |
| Ec WH59 | Antibiotic sensitive clinical<br>*E. coli* isolate | Westmead Hospital |
| Ec WH67 | Antibiotic sensitive clinical<br>*E. coli* isolate | Westmead Hospital |
| ***K. pneumoniae* strains** |  |  |
| Kp ATCC13883 | *K. pneumoniae* ATCC13883 | American Type Culture Collection (ATCC) |
| Kp WH49 | Antibiotic sensitive clinical<br>*K. pneumoniae* isolate | Westmead Hospital |
| Kp WH81 | Antibiotic sensitive clinical<br>*K. pneumoniae* isolate | Westmead Hospital |
| Kp WH84 | Antibiotic sensitive clinical<br>*K. pneumoniae* isolate | Westmead Hospital |

**Table 3.**
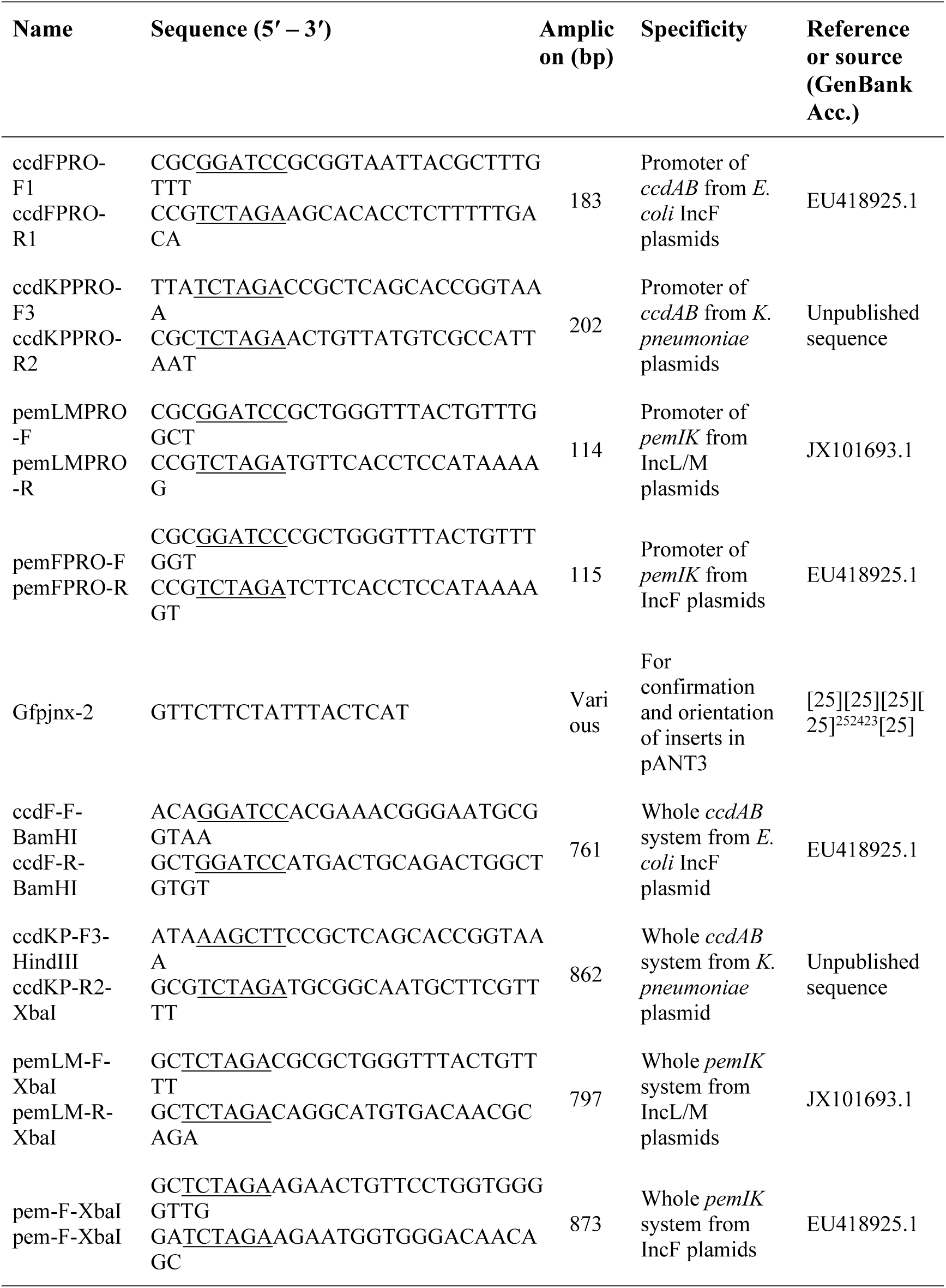
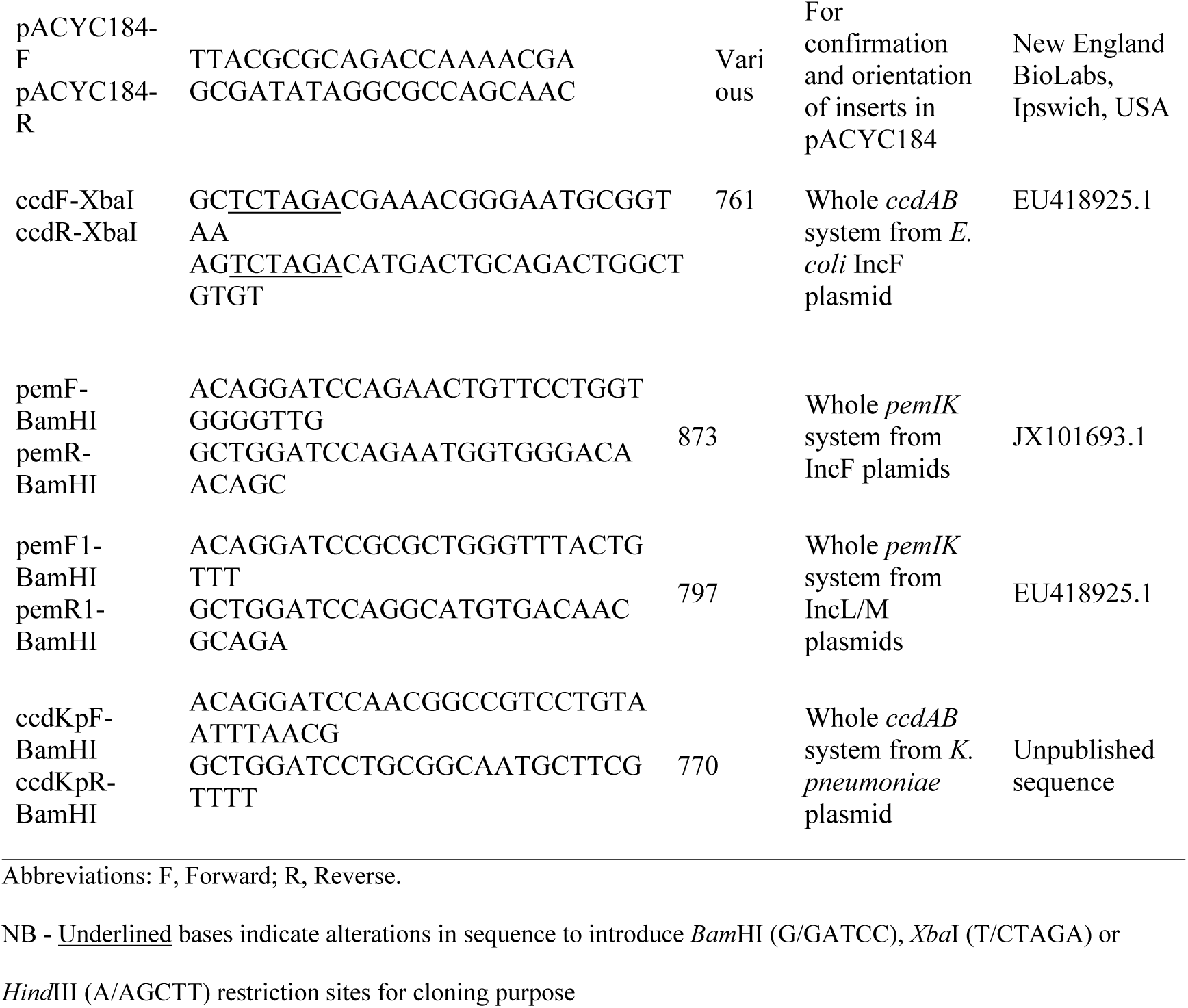
Oligonucleotide primers used in this study.

### Measurement of relative promoter strength

Relative strengths of putative TAS promoters were determined from TA promoter-*gfp* constructs using methods described previously [26]. Briefly, the predicted TAS promoters were cloned upstream of a promoterless *gfp* in the expression vector pANT3, and transformed into four strains each of *E. coli* and *K. pneumoniae*. Overnight cultures were inoculated from single colonies into LB broth with kanamycin, and grown with shaking at 37 °C. The cultures were then diluted 200 x in LB broth and grown under the same conditions for a further 3-4 h. Cells were harvested by centrifugation and the pellets resuspended in 0.85% sodium chloride (saline). The concentrations of each sample were then standardised to 1.5×10^8^ cfu/mL, as determined using the DensiCHEK™ Plus nephelometer (bioMérieux, France). Fluorescence was analysed using a Victor3 plate reader (Perkin Elmer, MA, USA), with an excitation wavelength of 485 nm and an emission wavelength of 535 nm. Experiments were performed in triplicate, and readings were averaged and corrected for background fluorescence by subtracting the pANT3 (no promoter) reading from each sample.

### Plasmid stability assays

To assess plasmid stability, the whole TAS (including the putative promoter and ribosome binding site) was cloned into a low and a high copy plasmid (pACYC184 and pBSCK+ respectively) and the constructs transformed into two strains each of *E. coli* and *K. pneumoniae*. Plasmid stability was assessed as described previously [27]. Briefly, a single colony of *E. coli* or *K. pneumoniae* bacteria carrying the relevant plasmid was grown in LB broth at 37 °C with shaking at 225 rpm without antibiotic. Bacterial cultures were transferred into fresh LB medium at 1:1000 dilution twice daily for 3 days (4 days for low copy plasmid constructs). Samples were taken before every transfer, diluted in saline and plated on to LB agar without antibiotic and incubated at 37 °C for 18 h. From each plate, 120 colonies were replica plated onto LB agar plates with and without the indicator antibiotics to estimate plasmid retention.

## Results and Discussion

### Distribution of type II TAS in the plasmids found in *K. pneumoniae* strains

Twenty-seven different putative type II TAS were identified (Table 4), with *ccdAB, pemIK* and *vagCD* most common among them (S1 Table). We compared the distribution of the TAS identified here in *K. pneumoniae* plasmids with previously reported TAS in *E. coli* plasmids [11]. *E. coli* and *K. pneumoniae* are closely related members of the *Enterobacteriaceae* family, sharing a large number of mobile antibiotic resistance genes, mainly via plasmids. However, only three TAS (*ccdAB, pemIK*, and *vagCD*) were common in both, with *ccdAB* and *pemIK* most commonly shared. Many, such as *MNT-HEPN*-like, *GNAT-RHH*-like, *Bro-Abr-*like and *Bro-ArsR-*like TAS appear to be most important in *Klebsiella* plasmids and have not previously been reported in *E. coli* plasmids [11, 13].

**Table 4.** Summary of TAS found on *K. pneumoniae* plasmids.

| Plasmid type | No. of plasmid | Type II TA Systems* |
| --- | --- | --- |
| IncF | 568 | None**, *vagCD*, *higBA*, *ccdAB*, *pemIK*, *GNAT-RHH* like, *MNT-HEPN* like, *COG5654-5642*, *relE-PhD* like, *relE-Xre* like, *pfam13420-TIGR01764*, *mazEF*, *mazF-RHH* like, *vapBC*, *parE-relB*, *relBE*, *MNT-RHH* like, *hicAB*, *fic-PhD* like, *relE-COG2442* like, *relE-COG5606* like, *COG12446-pfam13384* |
| IncR | 76 | *pfam13420-TIGR01764*, *vagCD*, *ccdAB*, *MNT-HEPN* like, *relE-Xre* like, *mazEF*, *mazF-AbrB* like, *relBE*, *pemIK*, *higBA*, *mazF-RHH* like, *GNAT-RHH* like, *COG5654-COG5642* |
| IncA/C | 62 | None, *pemIK*, *relE-Xre* like, *pfam12658-pfam00126*, *vagCD*, *GNAT-RHH* like |
| IncX | 60 | None, *hicAB*, *relE-Xre* like, *pfam13420-TIGR01764*, *vagCD*, *relBE* |
| IncH | 58 | *hipA-Xre* like, *relBE*; *COG5654-COG5642*; *GNAT-RHH* like, *relE-PhD* like, *relE-Xre* like, *MNT-HEPN* like, *vagCD*, *pemIK*, *pfam13420-TIGR01764*, *pfam12568-pfam00126* |
| IncL/M | 51 | None, *pemIK*, *pfam13420-TIGR01764* |
| IncN | 33 | None, *vagCD*, *MNT-HEPN* like |
| IncI | 10 | None, *hicAB*, *pfam13420-TIGR01764* |
| IncQ | 7 | *higBA*, *mazF-RHH* like, *pfam12568-pfam00126*, *relE-Xre* like, *GNAT-RHH* like, *pemIK*, *vagCD*, *COG12446-pfam133844* |
| IncU | 2 | None |
| IncW | 1 | None |
| Not determined | 114 | None, *relE-Xre* like, *GNAT-RHH* like, *higBA*, *mazF-RHH* like, *MNT-HEPN* like, *vagCD*, *relBE*, *Bro-Abr* like, *Bro-ArsR* like, *COG5654-COG5642*, *pfam13420-TIGR01764*, *higBA*, *mosTA*, *pfam00583-pfam00376* |

\* The TAS listed are all the TAS found on plasmids of that Inc type. Individual plasmids may have none, one or a combination of the listed TAS. For a more detailed distribution of the TAS on each individual plasmid, please see Supplementary S1 Table.
\*\* None = no type II TAS found on the plasmid.

Plasmids found in *K. pneumoniae* species belonged to 11 different Inc types (Table 4), although around 10% (114/1063) were not assigned an Inc type in PlasmidFinder. More than half of the plasmids were from the IncF replicon group, which is the most common plasmid replicon type in *Enterobacteriaceae* [28]. Replication gene variation can be used to further subdivide IncF plasmids (e.g. IncFIA, IncFIB, IncFIC, IncFII etc) and IncFII plasmids are usually further subdivided with a subscript to indicate the typical host species, e.g. IncFII_K_ and IncFII_S_ for the IncFII plasmids found in *Klebsiella* and *Salmonella* species, respectively. For simplicity, all IncF plasmid subtypes found in *K. pneumoniae* strains were grouped as IncF in this study. A number of *K. pneumoniae* plasmid types (L, M, X, A, C, N and H) (Table 4) are also commonly found in *E. coli* and other *Enterobacteriaceae*, [29] but some that are quite commonly reported in *K. pneumoniae* (e.g. IncR, representing 76/1083 plasmids) are rarely reported in *E. coli*.

Other associations are noteworthy. For example, *ccdAB* is somewhat specific to IncF plasmids in *K. pneumoniae*, while *pemIK* is common to both IncL/M and IncF plasmids (Table 4, S1 Table), in line with previous observations [9]. *vagCD* is another TAS predominantly found on the *bla*_CTX-M_-carrying IncF plasmids in *E. coli* (also as previously noted) [11], and in *K. pneumoniae* (Table 4). However, *vagCD* was also common in IncR in *K. pneumoniae* (35 of 76) but not in *E. coli*. Most IncX plasmids lacked an easily identifiable TAS, save for a few carrying *hicBA*, also as previously noted [30].

### *ccdAB* sequences differ between species, while *pemIK* is highly conserved

The two predominant TAS modules, *ccdAB* and *pemIK*, across both *E. coli* and *K. pneumoniae* plasmids (Table 4, [11]) were compared. Alignment of *ccdAB* from IncF plasmids derived from two species (*E. coli* and *K. pneumoniae*; henceforth termed “*ccdAB*_*EC*_” and “*ccdAB*_*KP*_” respectively) revealed 85% identity in the nucleotide coding sequence (S1A Fig), translating to 75% and 93% amino acid sequence identities in the antitoxin and toxin respectively (Fig 1A). Secondary structure analysis revealed that although the antitoxin ccdA had the same basic structure in both cases, the toxin ccdB derived from *K. pneumoniae* had an additional small α-helix near the C-terminal end (S2 Fig). The putative promoters were less similar, with only a 33% identity throughout the promoter region (−10, −35 and spacers) (Fig 1C). Thus, it is expected the expression of these TA modules in different species would vary.

**Fig 1.**
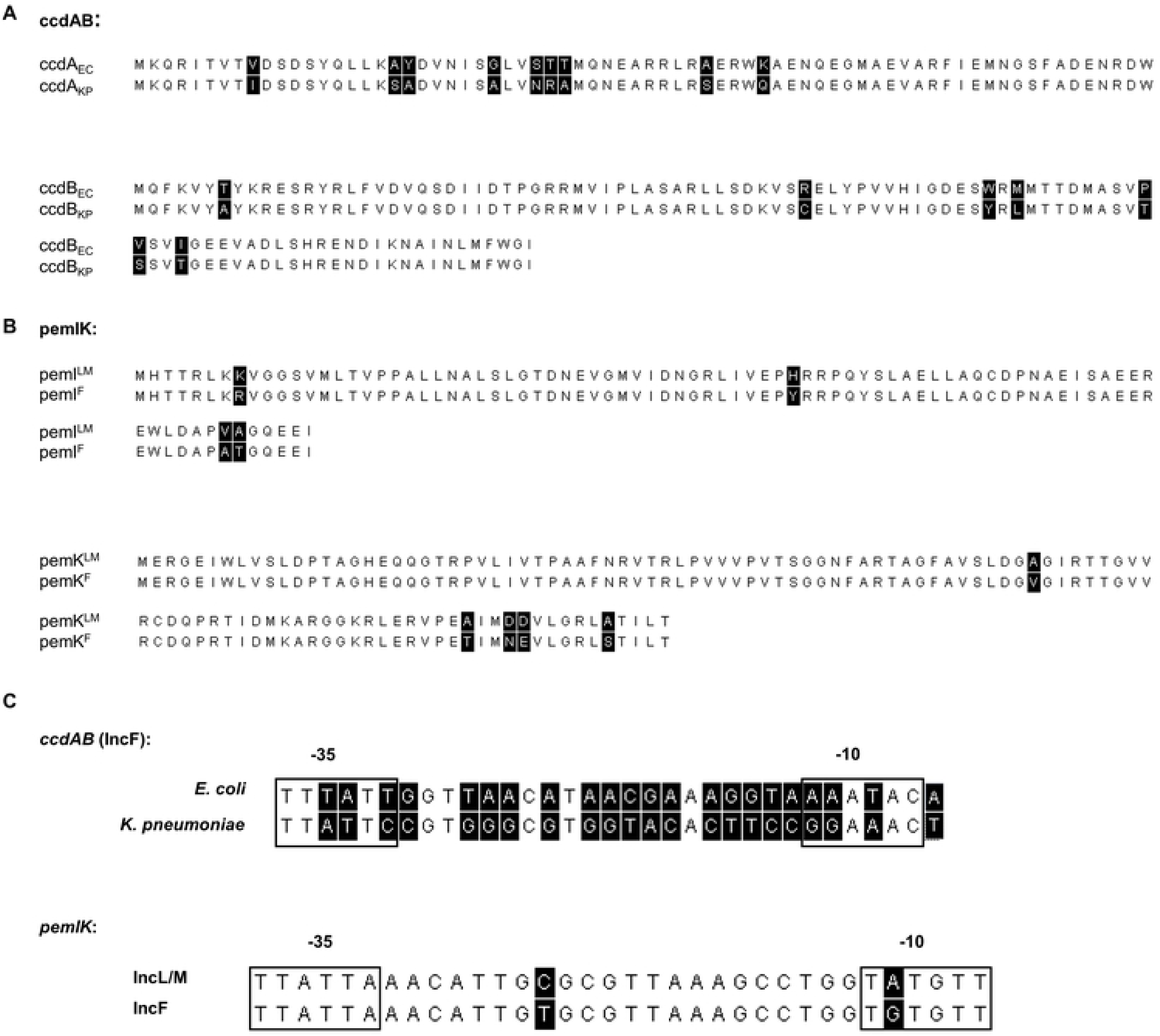
Sequence alignments of *ccdAB* and *pemIK*. These include the amino acid sequence alignments of (**A**) the antitoxin ccdA and the toxin ccdB, from *E. coli* and *K. pneumoniae*; and (**B**) the antitoxin pemI and the toxin pemK, from IncL/M and IncF plasmids; and (**C**) the nucleotide alignments of the putative promoter regions of the selected TAS. The putative −35 and −10 elements are boxed. All sequences were aligned in MEGA7 using the ClustalW algorithm. Non-identical residues/bases are highlighted in black.

By contrast, *pemIK* sequences differed with the plasmid Inc types, rather than with the host bacterial species. Alignment of *pemIK* from two different plasmid types, IncL/M and IncF (henceforth “*pemIK*^*LM*^” and “*pemIK*^*F*^” respectively) revealed 92% identity in the nucleotide coding sequence (S1B Fig), translating to 95% amino acid sequence identities in both proteins (Fig 1B). The pemI antitoxins and pemK toxins also had identical secondary structures (S3 Fig). Similarly, the putative promoters had 94% nucleotide identity (Fig 1C) with only one SNP in the −10 sequences, indicating that *pemIK* TAS is quite conserved within different plasmid types.

### Variation in expression of *ccdAB* in different host strains is greater than expression of *pemIK*

GFP expression from both *pemIK* promoter variants were relatively strong in all strains/species tested, with expression higher than from either of the *ccdAB* promoters tested, as well as the workhorse *tac* promoter [31] (Fig 2), in line with the apparently minor variation in *pemIK* promoters. *pemIK* is the sole TAS evident in published IncL/M plasmids [9], and these data suggest that *pemIK* alone is sufficient to provide stability to the plasmids carrying them, regardless of the host bacterial species.

**Fig 2.**
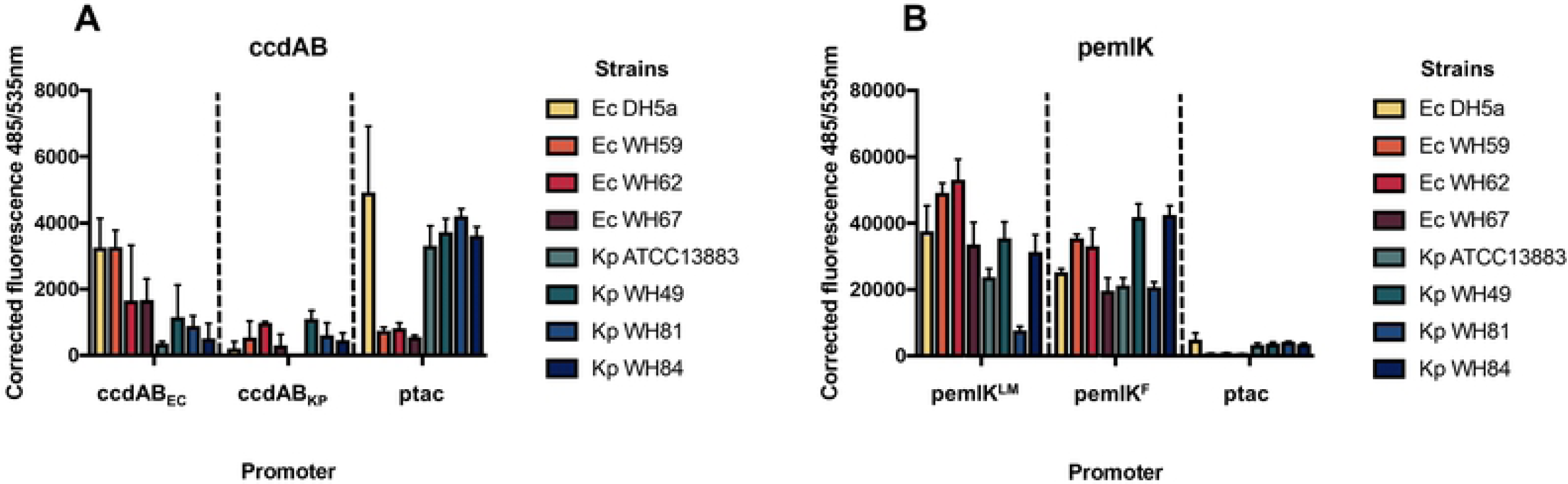
Relative expression of GFP from promoters of (A) *ccdAB* and (B) *pemIK*. The putative promoter of each TAS was inserted upstream of promoterless *gfp* in pANT3 and transformed into four *E. coli* and four *K. pneumoniae* strains. pANT5, with the *tac* promoter (ptac)-*gfp* construct, served as a positive control, and values are corrected for background noise. Data shown are the means of three replicates, with the error bars representing one standard deviation from the mean. Please note the ten-fold differences in y-axis scales.

In contrast, expression of GFP from *ccdAB* promoters is generally lower than that of *pemIK* and showed host species specificity. GFP expression from the *E. coli* specific *ccdAB* promoter was higher in *E. coli* than in *K. pneumoniae* strains. The expression of GFP from the *K. pneumoniae* specific *ccdAB* promoter also varied between strains in each species but was relatively lower than from all other promoters tested here.

### *ccdAB* function is specific to host strain, whereas *pemIK* is not

To assess whether the apparent specialisations within *ccdAB* (by host species) and *pemIK* (by plasmid type) are also reflected functionally, we assessed plasmid stability in two host strains of each species, in both low and high copy number plasmids. The low copy plasmid data are presented (Fig 3) as the natural plasmids carrying these TAS are generally low copy number. Consistent data for high copy plasmids can be found in the supplementary material (S4 Fig).

**Fig 3.**
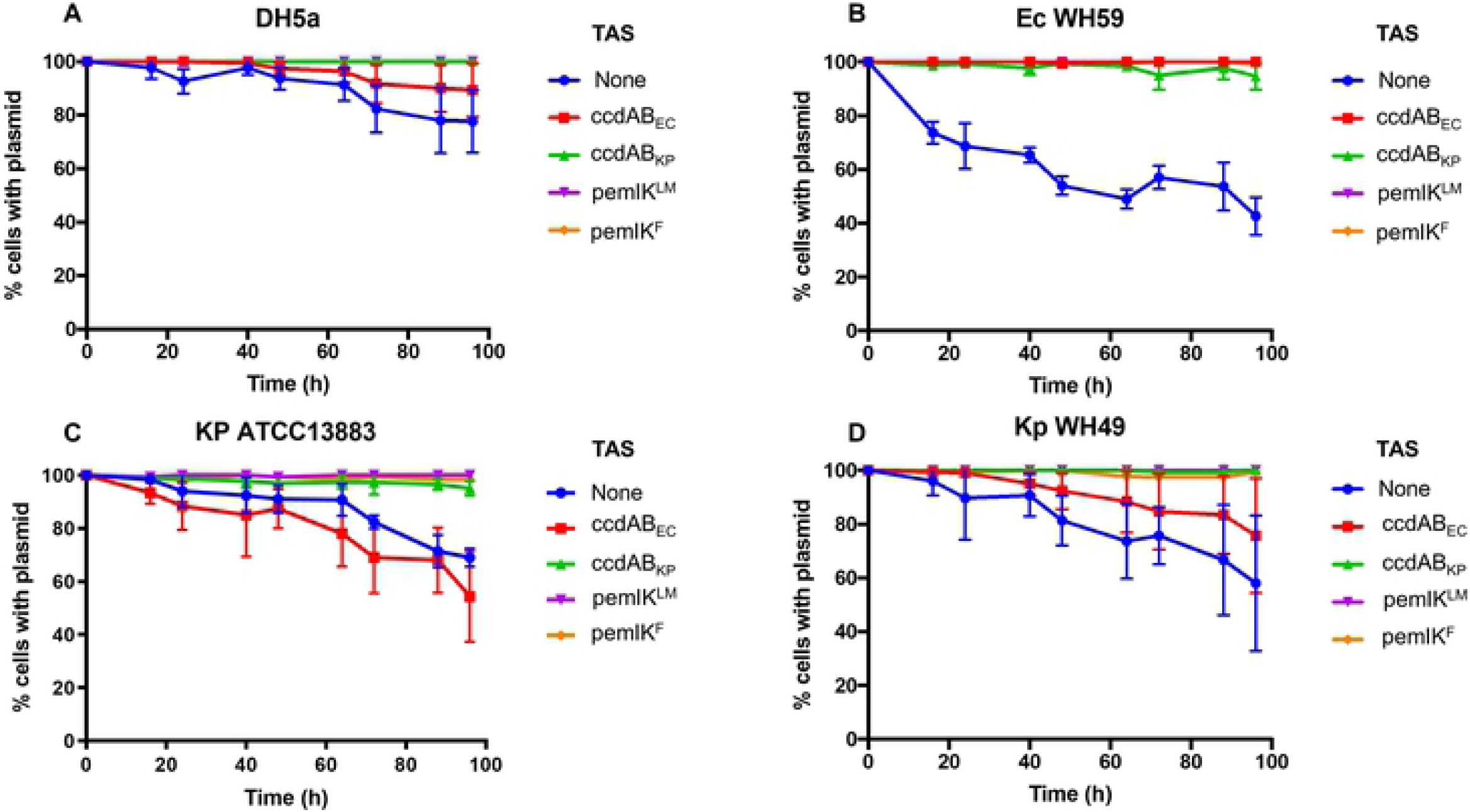
Stability of a low copy number plasmid with and without the specified TAS over 96 h. Each TAS was cloned into a pACYC184 backbone, and the percentage of cells retaining the plasmid calculated at each time point. Measurements were done in two species, *E. coli* (**A**: DH5α and **B**: Ec WH59) and *K. pneumoniae* (**C**: Kp ATCC13883 and **D**: Kp WH49). Data shown are the means of three replicates, with the error bars indicating one standard deviation from the mean.

The plasmid stability conferred by *ccdAB* variants varies between host species (Fig 3) and is generally consistent with the GFP expression data, with *ccdAB*_*EC*_ stabilising plasmids in both *E. coli* strains but not in *K. pneumoniae* strains. *ccdAB*_*KP*_, on the other hand, appears to be less specialised, conferring some plasmid stability in all strains, regardless of species. Given the relatively low level of expression from *ccdAB* promoters and discrepancies in the conferred plasmid stability across species and strain, it is perhaps unsurprising that it is typically one of several TAS in IncF plasmids [11]. By contrast, the more reliably and vigorously expressed *pemIK* variants conferred significant plasmid stability in all strains tested (Fig 3).

Overall, *pemIK* appears less specialised than *ccdAB*. The sequences and expression of *pemIK* variants differed relatively less, and plasmid stability functions of both variants were more consistent. In contrast, *ccdAB* appears to be less widely distributed and more variable in sequence and function, different variants conferring plasmid stability in different strains.

The inability of *ccdAB* to confer significant plasmid stability in certain host strains raises the possibility that it is performing another role in these strains. Previous work has demonstrated the involvement of both plasmid and chromosomally encoded TAS in a range of functions, including persister cell formation, virulence, antibiotic and heat tolerance, and other bacterial stress responses. [17, 27, 32-35] This suggests that some TA systems have specialised functions other than plasmid maintenance and that these may contribute to the epidemiology of the conjugative plasmids which carry them.

## Acknowledgements

Thanks to Sanaz Pakbaten Toupkanlou for providing two plasmid constructs.

## Supporting Information

**S1 Table. Distribution of type II TA systems in the plasmids of *K. pneumoniae*.** (XSL file).

**S1 Fig. The alignment and comparison of the nucleotide sequences of *ccdAB* and *pemIK***. Nucleotide sequence alignments of the coding region of (**A**) *ccdAB* from plasmids found in *E. coli* and *K. pneumoniae*, and (**B**) *pemIK* from IncL/M and IncF plasmids. Sequences were aligned in MEGA7 using the ClustalW algorithm. Non-identical residues are highlighted in black.

**S2 Fig. Comparison of the predicted secondary structures of *ccdAB*.** The predicted secondary structure of the *ccdAB* encoded toxins and antitoxins from plasmids found in *E. coli* and *K. pneumoniae*. Arrows represent β-strands, cylinder shapes represent α-helices and lines represent random coils. Pred: predicted secondary structure; AA: amino acids; numbers below each structure represent the amino acid positions within the proteins.

**S3 Fig. Comparison of the predicted secondary structures of *pemIK*.** The predicted secondary structure of the *pemIK* encoded toxins and antitoxins from IncL/M and IncF type plasmids. Arrows represent β-strands, cylinder shapes represent α-helices and lines represent random coils. Pred: predicted secondary structure; AA: amino acids; numbers below each structure represent the amino acid positions within the proteins

**S4 Fig. Plasmid stability effects of *ccdAB* and *pemIK* in a high copy plasmid.** Stability of a high copy number plasmid with and without TAS over 72 h. Each TAS was cloned into a pBCSK+ backbone, and the percentage of cells retaining the plasmid calculated at each time point. Measurements were done in two species, *E. coli* (**A**: DH5α and **B**: Ec WH59) and *K. pneumoniae* (**C**: Kp ATCC13883 and **D**: Kp WH49). Data shown are the means of three replicates, with the error bars indicating one standard deviation from the mean.

